# Estimating dead fish quantities dropping out of gillnets when direct observations are impossible

**DOI:** 10.1101/2025.04.23.650286

**Authors:** Hugues P. Benoît, Jean-Martin Chamberland, Jacques Allard

## Abstract

Fish that are caught in passive or fixed fishing gear and subsequently die and drop out of the gear, or that are removed by scavengers, do not show up in catch statistics and constitute unaccounted-for removals. Estimating these removals is challenging, particularly when direct observations are not possible because fishing conditions preclude the use of cameras or other devices or means to catch or observe the removals. Using a simple theoretical process model, we define two independent modelling approaches to estimate unaccounted mortalities resulting from drop-out. The first is based on a minimally refined analysis of commonly available fishery catch per unit effort data. The second, models data that can be obtained straightforwardly from field experiments and from at-sea fishery observers, specifically data on catch amounts, and catch composition according to three condition categories – live, dead-fresh and degraded. We apply these modelling approaches to data from the Gulf of St. Lawrence (Canada) Greenland halibut (*Reinhardtius hippoglossoides*) gillnet fishery, for which multi-day soak durations have long been suspected of generating significant unaccounted fish death while only little evidence is available. In fact, observed proportions of dead decaying and ultimately discarded catch for this fishery are small and increase only a little with increased soak duration, constituting seemingly contradictory evidence. The two modelling approaches produce similar results, estimating that total dead catches of Greenland halibut were on average 4.7 to 5.4 times the recorded landings over the period from 2000 to 2024. This result is consistent with the high mortality levels suggested by the assessment for this stock. Of broader relevance to other gillnet fisheries worldwide that may employ shorter soak duration, we found that estimated unobserved catch losses equalled retained catch after soak durations of 15 hours or less. Importantly the condition of fish in catches may belie the quantities of dead fish losses. Failing to account for fishing derived loss of the magnitude estimated here could result in important biases in stock assessments and associated sustainable harvesting frameworks, in addition to constituting an important source of waste.

## 1. Introduction

Incomplete accounting of the amount of fish killed by fishing is likely to result in underestimation of stock productivity, and possibly in time-varying biases in abundance and fishing mortality estimated from assessments, particularly if the fraction of unaccounted catch changes over time (Punt et al. 2006, Dickey-Collas et al. 2007, Perretti et al. 2020). In turn, this will affect estimated reference points, and potentially the reliability of a precautionary approach framework for the stock. Catch misreporting and the discarding of dead fish by harvesters are commonly cited sources of under accounting of dead catch (Kelleher 2005, Bousquet et al. 2010, Cadigan 2016, Cook 2019), although many other sources exist, including the mortality of fish escaping gear and ghost fishing (ICES 2005, Uhlmann and Broadhurst 2015).

Fish that are caught in passive or fixed fishing gear and subsequently die and drop out of the gear, or that are removed by scavengers, constitute a potentially important source of unaccounted-for removals (Uhlmann and Broadhurst 2015). Quantifying these removals has been particularly challenging. Direct observation is not straightforward, and in many fisheries is simply impossible. Limited natural light and/or field of view preclude the use of cameras (e.g., Harrison and Harvey, 2009) and acoustic imagery. These factors, in addition to depth, preclude the use of divers, which, like submersibles, area also limited to episodic sampling. The use of artificial light risks interfering with capture and scavenger processes. It is also not feasible to accurately recover and count dead animals that dropped from nets and into a bottom purse net (sensu Hay et al. 1982) given that dropped out individuals are likely to be in an advanced state of decay and may be more likely to be depredated and removed or lost from the purse net. Finally, while depredation rates from gillnets can be estimated using marked fish that are meshed before net deployment (Glemarec et al. 2024), this approach is not suitable for estimating drop-out losses because the meshed fish will not correctly mimic the entire process of capture, mortality, degradation and loss. Drop-out losses must therefore be inferred when direct observations are not possible.

Gillnets are used exclusively in the directed commercial fishery for Greenland halibut (*Reinhardtius hippoglossoides*) in the Gulf of St. Lawrence (GSL, Canada), Northwest Atlantic Fisheries Organisation (NAFO) divisions 4RST. Historically, a large proportion of fishing trips have employed prolonged gillnet soak duration, often exceeding the regulated 72 hours maximum soak time (Figure 1; Chamberland and Benoît, 2024). These soak durations considerably exceed those employed in other groundfish fisheries in the area, notably cod (Figure 1), and are in line with values that have previously been associated in other areas with degradation of the catch to the point at which some may fall out of the gear or be depredated before the gear is retrieved (Ward et al. 2004, Patterson et al. 2017). Anecdotally, some GSL Greenland halibut harvesters speak of fishing their nets effectively as self-baited nets, implying a wilful practice of allowing some catch to decompose to increase fishing power. This behaviour is not unexpected, as explicitly baited gillnets have been shown to significantly increase Greenland halibut catch rates (Engås et al. 2000, Bayse and Grant 2020).

**Figure 1.**
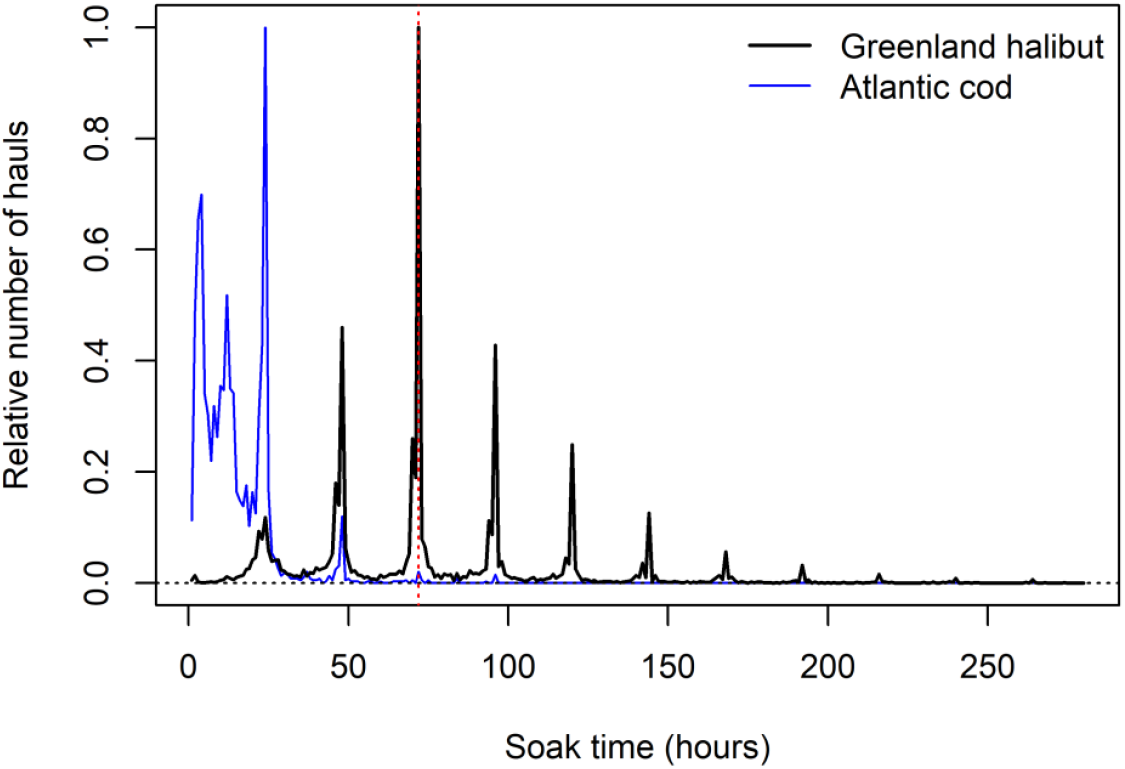
Relative frequency distribution of gillnet soak durations for Greenland halibut and Atlantic cod directed fisheries in the Gulf of St. Lawrence, 2000-2024. Durations were classified into one-hour bins and scaled to the maximum bin-specific value for each species. The dashed red vertical line indicates the regulated 72 hour maximum untended soak duration for Atlantic Canadian groundfish fisheries.

The assessment for GSL Greenland halibut indicates very elevated mortality for large individuals captured in the fishery since at least the 1990s (Chamberland and Benoît 2024). This is in contrast to what are otherwise low estimated exploitation rates based on reported landings, with values <10% in most years. Unaccounted-for removals may therefore explain much or all of the apparent excess mortality, for what should otherwise be a relatively long lived species (Brogan et al. 2021). Establishing the magnitude of mortality caused by drop-out is key to understanding how fishing affects the stock and for establishing reliable approaches for its sustainable management.

In the GSL Greenland halibut fishery, fish that are unsuitable for human consumption may be discarded. Data from at-sea observers show that the proportion, by weight, of the directed species catch that is discarded increases with soak duration, to a median value of about 9% for soak durations of 8 days or more (Figure S1). While these results indicate the potential for catch losses due to drop out, they assuredly underestimate the amounts, potentially considerably. Trivially, the percent losses will be underestimated because the weight of decayed remains will underrepresent the constituent number of fish, compared to the weight of retained catch. Importantly, the weight of degraded fish, a state variable, provides no information on the rate at which degraded fish mass turns over in the net.

In this paper we develop a simple theoretical process model, from which we define two independent modelling approaches to estimating unaccounted mortalities in this fishery. The framework relies on modelling the turnover of fish in gillnets over the duration of soaking. The first approach is based on a minimally refined analysis of fishery catch per unit effort data and models turnover implicitly. This approach is applicable to fisheries like GSL Greenland halibut that use catch degradation to as part of the fishing strategy. The second approach explicitly models turnover using data on catch amount and catch composition according to three condition categories – live, dead-fresh and degraded – from field experiments and from at-sea fishery observers. We apply these modelling approaches to the case of the GSL Greenland halibut gillnet fishery.

## 2. Methods

### 2.1 Data sources, including field experiments

The study involves data from four sources: fishery statistics, field experiments, a process study conducted during the field experiments, and data from at-sea observers.

Data from the commercial fishery were extracted from Fisheries and Oceans Canada’s Zonal Interchange File Format database, which compiles fishing-trip specific observations originating from different fishery monitoring programs, including dockside catch monitoring and logbooks. The data extracted were landed amount (kg), the number of gillnets used, the soak duration, and the date and the NAFO subarea for each fishing trip for the years 2000-2024.

Four separate field experiments were undertaken aboard chartered vessels, respectively in June and in July 2022, in May 2023 and in September 2024. Each experiment consisted of fishing gillnets using soak durations ranging from 2 to 126 hours, across 21 to 32 hauls. The total number of hauls and the soak durations varied somewhat among experiments due to constraints associated with weather and crew availability. Experimental sites were all in the St. Lawrence Estuary, but locations differed between experiments. Within experiments, nets were separated by distances comparable to those used in the fishery. The total number of Greenland halibut according to each of three condition categories was recorded for each haul:

> *Live* – the fish displays some movement, including in response to manipulation.
>
> *Dead-fresh* – the fish displays no response to manipulation, the flesh is firm.
>
> *Degraded (dead)* – flesh is very soft to the touch or only bones remain, parts may be missing.

During the field experiments, a process study was undertaken to estimate the rate of degradation of dead fish. Four live fish from a previous haul were measured and then attached to each gillnet, two on the headline and two on the leadline using a plasitic tie-wrap run through the mouth and operculum, along with a uniquely numbered rubber label. The gillnet was then soaked as part of the regular experiment, and the condition of each of the four fish upon retrieval was noted. Although these fish were alive when attached to the net, they are assumed to have died shortly after the net was deployed as a result of air exposure and handling before their return to the water, and constraints on breathing from the tie-wrap. The number of fish used in the process study was kept as small as feasible, noting that these fish would otherwise have been retained and landed, and therefore killed nonetheless.

At-sea observers are deployed in the directed fishery at a minimum target rate of 5% of fishing trips to monitor regulatory compliance, the size and sex composition of target species catch, and the bycatch. In 2021-2023, observers also collected, for the purposes of the present study, data on the condition of Greenland halibut captured, using the condition categories defined above. From two selected hauls per day, observers collected condition observations for 25 individual fish, often the first fish brought aboard during the haul. Those observations were associated to data on the characteristics of the haul (e.g., date, location, soak duration) which are collected by observers as part of their routine sampling.

### 2.2 Modelling – General framework

Let *N*(*t*) be the new instantaneous catch in the gillnet at soak time *t*, where *t*=0 when the gillnet is deployed. *N*(*t*) is unknown in the present application, and would otherwise be very difficult to observe in most fishery contexts.

Let *L*(*t*) be the catch available and of sufficient quality to land at time *t*, hereafter landable catch. *L*(*t*) is known from observations, either landings in fishery catch statistics or the catches during experiments. *L*(*t*) excludes catches that are discarded at sea because they are decomposed or degraded and therefore unmarketable.

*S*(*u*) is a retention function, i.e., proportion of catch that remains in the net and is of sufficient quality to be landed for a time *≥ u* after capture; *S*(0) = 1, *S*(*∞*) = 0.

The retention function is central to the estimation of catch loss as it describes the rate at which fish drop out of the net or become too decomposed to land and sell. In this paper, we present two general and independent approaches to its estimation (details below).

The process that generates available catch is defined as:

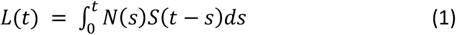

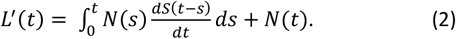

An exponential retention function provides a simple description of a process that involves a fish’s death and subsequent decomposition following capture, although other functions are possible (see below):

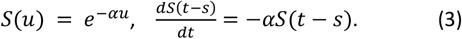

Substituting eq 3 into eq 2:

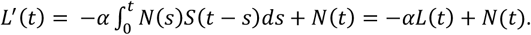

Solving for *N*(*t*), we obtain:

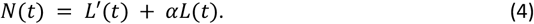

For an arbitrary survival function *S*(*u*), *N*(*t*) can be computed numerically from *L*(*t*) by discretizing equation (1), as follows.

*i* = 0 … *I*

*t*_*i*_ and *u*_*i*_ are time sequences, *t*_0_ = *u*_0_ = 0

*L*(*t*_*i*_) = (*I +* 1) *×* 1 column vector of landings at time *t*_*i*_

*N*(*t*_*i*_) = (*I +* 1) *×* 1 column vector of new catch at time *t*_*i*_

*S*(*u*_*i*_) = Retention function, recalling *S*(0) = 1.

From *S*(*u*_*i*_), the following square transition matrix is created:

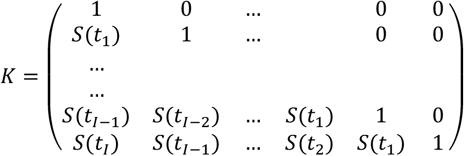

Discretizing eq 1 gives

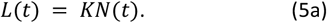

Given that the determinant of *K* is 1, *K* can be inverted and therefore,

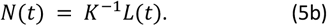

The extent to which landings made using a soak duration of *t*_*i*_ underrepresent the total number of fish killed as a result of capture to *t*_*i*_ can be expressed using the following dead-catch ratio function:

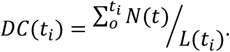

In turn, an average dead-catch ratio for year *y* is calculated as:

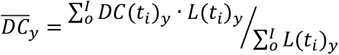

where *L*(*t*_*i*_)_*y*_ denotes the total landings in the fishery in year y associated with a soak duration of *t*_*i*_.

#### 2.2.1 Model 1 – Estimation from fishery catch-per-unit-effort

For simplicity, catches in the GSL Greenland halibut fishery are hypothesized to originate from two processes. Fish that are in the area where a gillnet is set are assumed to be captured rapidly, resulting in an initial capture and potential landing, *L*_*local*_. These fish die, decay and drop out of the net according to an exponential retention function, exp(−*αt*). The decaying fish attract fish from the broader regional area, that, in turn, are captured, die and drop out, perpetuating the attraction and capture process. Formally, the entire process is expressed as:

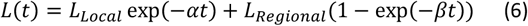

where,

*L*_*Regional*_ is the density of fish in the broad regional area, which is assumed to not be depleted by capture at the gillnet site during the soak period; and

(1 − exp(−*βt*)) describes a process of initially increased attraction to the gillnet location, eventually reaching an equilibrium with the retention rate.

Eq. (6) describes a landable catch series that starts at the level *L*_*local*_, decreases initially with soak time, before increasing again to reach an asymptote at *L*_*Regional*_ (Figure 2, top panel). The form of the predicted *L*(*t*) series matches that of average standardized landings as a function of soak time in the Greenland halibut fishery estimated by Chamberland and Benoît (2024; Figure 3). Fitting to those standardized landings is the basis for the estimation of the parameters of eq. 6, and the prediction of *N*(*t*) and *DC*(*t*), as detailed below. The *N*(*t*), calculated using eq 5b and the *L*(*t*) from eq 6, comprises new catches that drop rapidly from a high level at a soak time of zero to rapidly reach a minimum level after a few hours, before abruptly reversing direction, to increase gradually to an asymptote (Figure 2, bottom panel).

**Figure 2.**
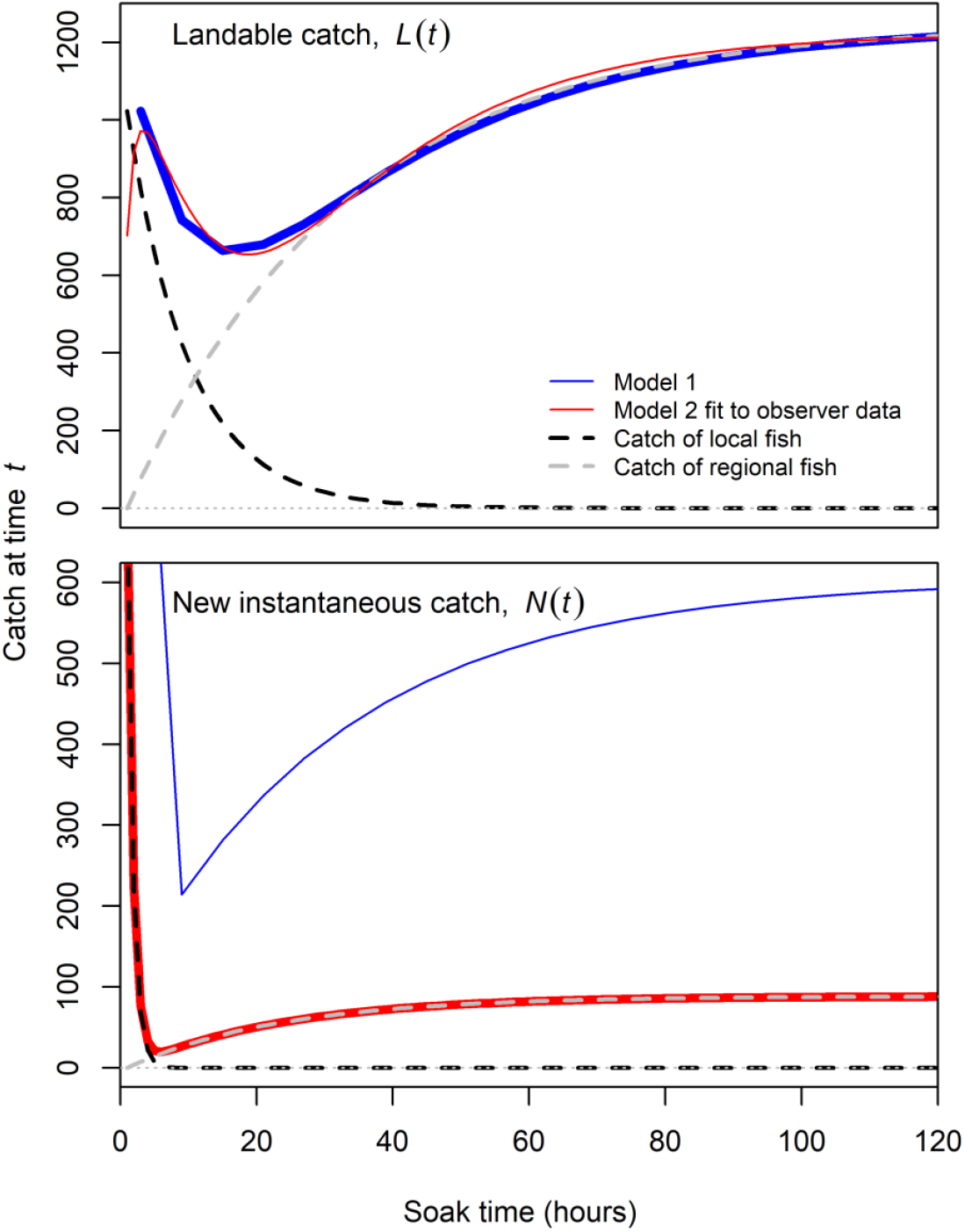
Schematic of the time series for landable catch (*L*(*t*); top panel), and new instantaneous catch (*N*(*t*); bottom panel), and how these are obtained for each of the two modelling approaches. In Model 1, *L*(*t*) (thick blue line) is modelled as comprising a decreasing function of landings of local fish over time (dashed black line) and an increasing asymptotic function of landings of regional fish attracted to the fishing site (dashed grey line; top panel); meanwhile *N*(*t*) is inferred using eq. 5b (thin blue line, bottom panel). Model 2 involves modelling *N*(*t*) as a latent process (thick red line; bottom panel) resulting from the capture of local fish and regional fish attracted to the area, and explicit modelling of the retention function *S*(*u*), and application of eq. 5a are used to predict *L*(*t*) (thin red line, top panel). Results for Model 2 in the figure are those obtained by fitting to all data, including those from at-sea observers, and provide an example of predictions from that modelling approach.

**Figure 3.**
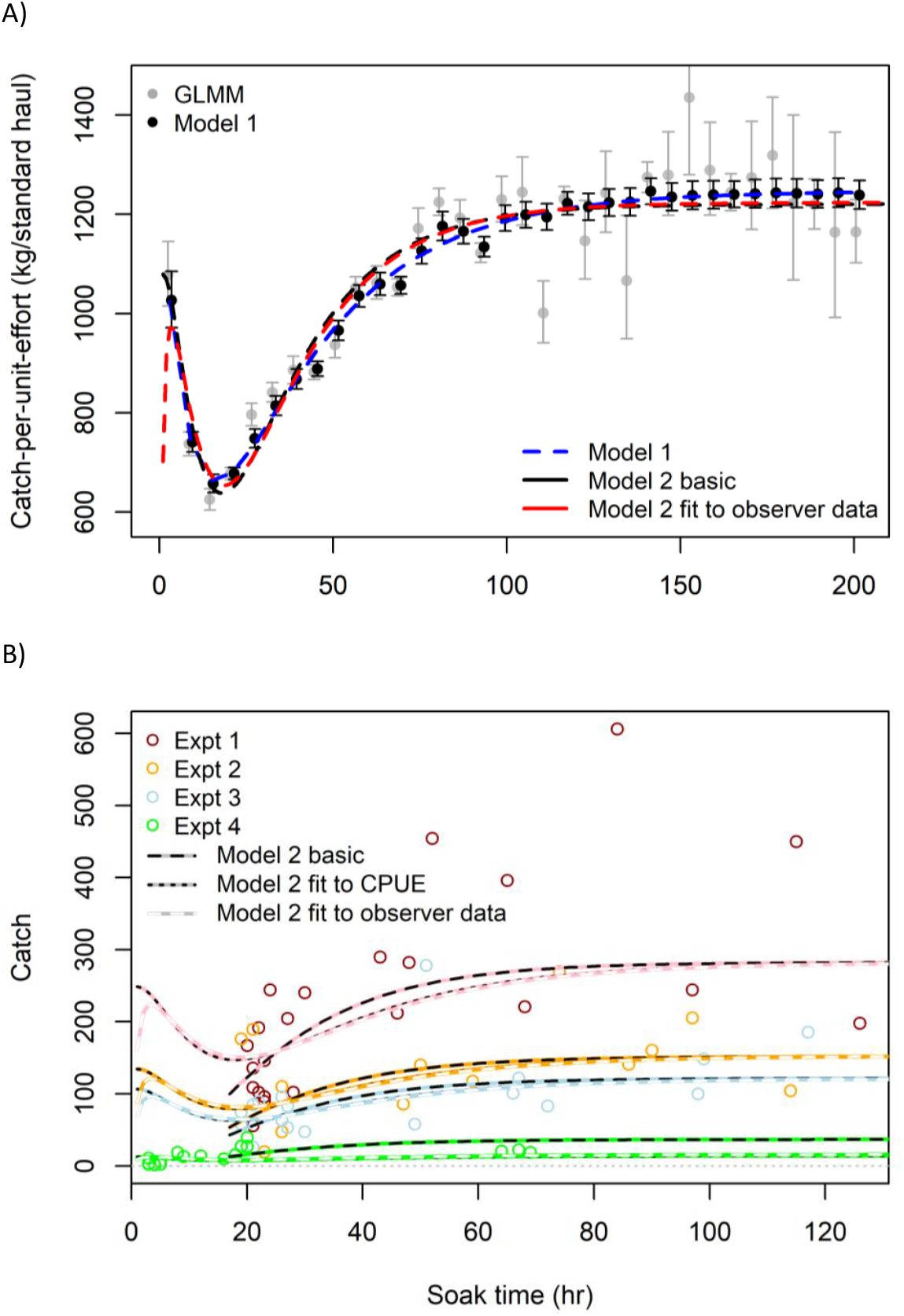
A) Standardized catch rate with standard error as a function of soak duration, estimated independently in a stand-alone generalized linear mixed-effects model (GLMM grey circles and bars) and estimated as part of fitting Model 1 (black circles and bars), along with model predictions for landable catch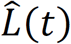, for Model 1 (dashed blue line), Model 2 fit to just the data from the experiments (black line) and Model 2 fit to all data, including from at-sea observers (red line). B) Fits of the three variants of Model (distinguished by line type) to the catch amounts in each of the four experiments (distinguished by colour).

Alternative forms for the catch process in eq. 6 are possible; however the resolution of the data available for fitting Model 1 could only support the simpler assumptions made here. For instance, the contribution of local catch could be assumed to initially increase as local fish encounter the net, before declining as free-swimming local fish are depleted and catches degrade and drop out. Alternative shapes of the retention function are also possible, for instance, ones akin to Deevey’s well known Type I and Type III survival curves (Deevey 1947; the exponential function in eq.6 is analogous to Type II survival).

#### 2.2.2 Model 2 – Estimation from data on the condition and degradation rate of catches

Model 2 takes an inverse modelling approach for the estimation of catch loss amounts by explicitly modelling both the capture process, described by *N*(*t*), and the catch retention process that generates *S*(*u*_*i*_). Landings are then predicted using eq. 5a, and fitted to observations.

New catches comprise the capture of local fish and the capture of fish attracted to the fishing site as depicted in Figure 2 (bottom panel) and described by:

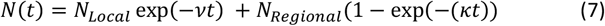

where *N*_*Local*_ is the density of fish at the fishing site, *ν* is the rate at which those fish are depleted through capture and *N*_*Regional*_ is the density of fish in the broader region. The term (1 − exp(−(*ϰt*)) conceptually describes the rate at which fish are attracted to the site, increasing rapidly initially as an increasing amount of gillnet captured fish decay and produce a bait plume, before reaching an asymptote when the smell of bait is diffused throughout the region.

Eqs 6 and 7 are clearly identical in form, although their conceptual derivation differs and they describe different properties of the catch, respectively landable catches and (latent) new catches. Importantly, the estimates of *L*(*t*) from Models 1 and 2 can be nearly identical across all soak time (see Results) or all but the shortest soak times (Figure 2, top panel shows an example of one implementation). Similarly, the two models can produce nearly identical trends in *N*(*t*) (Figure 2, bottom panel).

The catch retention process is modelled according to a series of three consecutive non-overlapping processes that initiates the moment a fish is caught, alive, in the gillnet. This fish eventually dies, once dead it begins to rot and degrade, eventually reaching a state in which it is no longer retained in the net and therefore drops out. Individually, the rates of mortality (*Z*_*M*_), degradation (*Z*_*F*_) and drop-out (*Z*_*D*_) were assumed to follow simple exponential decay functions, which seem appropriate for such processes. The entire process generates four condition classes of fish, whose relative composition in the catch as a function of the time following capture *u*_*i*_ is defined as:

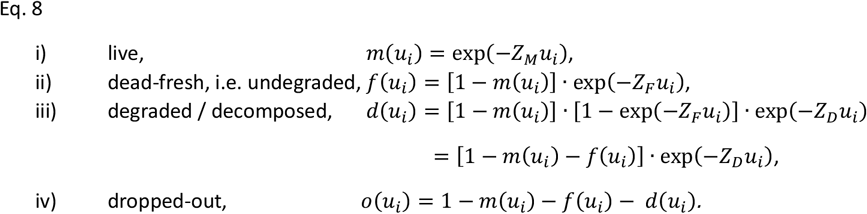

Preliminary analyses indicated evidence that some degraded fish may be sufficiently well meshed that they are unlikely to drop from the net even when only bones remain. To account for this possibility, we generalized eq 8iii to include a fraction of long-term ‘hangers-on’, *p*_*d*_, borrowing from the approach used in common cure-rate survival modelling (e.g., Farewell 1982):

iii) degraded / decomposed,

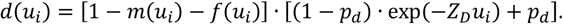

For each of these four classes, we constructed transition matrices, using the approach described earlier: *K*_*m*_, *K*_*f*_, *K*_*d*_, and *K*_*o*_. We then then estimated time series vectors for the number of fish in each category:

live, *M*(*t*) = *K*_*m*_*N*(*t*); dead-fresh, *F*(*t*) = *K*_*f*_*N*(*t*); degraded, *D*(*t*) = *K*_*d*_*N*(*t*); and, dropped-out, *O*(*t*) = *K*_*o*_*N*(*t*).

Based on the definitions provided earlier, *L*(*t*) = *M*(*t*) *+ F*(*t*), whereas *D*(*t*) + *O*(*t*) constitute unaccounted catch. The predicted values for *L*(*t*) can therefore be fitted to data on landings or experimental catches as part of the parameter estimation process.

Three of the condition classes are readily observable in gillnet catches: live, dead-fresh and degraded. The proportion of fish in condition class *c* in the gillnet, as a function of soak duration, *p*_*c*_(*t*), is simply,

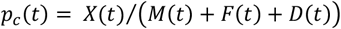

The predicted values for *p*_*c*_(*t*) can be fitted to observations of the relative (condition) composition of catches for individual gillnet hauls.

The parameters in Model 2 are not identifiable by simply fitting to data on catch amounts and catch composition. At least one of the rates involved in the retention process needs to be specified. Both the rates of mortality, *Z*_*M*_, and drop-out, *Z*_*D*_, are challenging to quantify accurately from observations. In contrast, the rate at which recently deceased fish decompose to the point of being unmarketable, *Z*_*F*_, can readily be estimated from observations from a process study such as the one described earlier.

### 2.3 Model fitting

All model fitting was undertaken in R, using the RTMB package v.1.6 (Kristensen 2024) to calculate the marginal negative loglikelihood for the models and the base R *nlminb()* function to estimate parameters.

#### 2.3.1 Model 1

The basic datum for fitting Model 1 was an individual landing, in kg, associated with a year, month and location of capture, the number of gillnets used and a gillnet soak duration (in hours). Location was classified by NAFO sub-area, while soak time was discretized into six-hour categories beginning with]0, 6]. Landings were standardized for differences in local density at the time of capture using generalized linear modelling (GLMM), much like in common fishery catch rate standardization (Maunder and Punt 2004). In contrast to common practice, standardized catches were estimated as a function of soak time category, rather than year. Details on the original GLMM fitting, model selection and validation are available in Chamberland and Benoît (2024).

Model 1 was fitted to the same input data and used the same GLMM structure as Chamberland and Benoît (2024). Specifically, individual fishery landings *Y*_*j,n,ym*_ associated with fishing trip *j*, in NAFO subunit *n*, and year and month *ym* were modelled as Tweedie (TW) random variables:

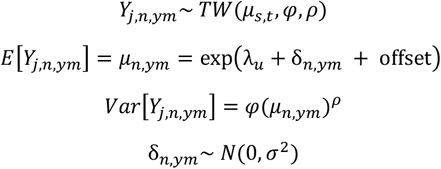

where,

*μ*_*n,ym*_ is the mean landed biomass in *n* and *ym*,

offset is the log of the number of gillnets

*φ* is the dispersion parameter of the Tweedie distribution (Dunn and Smyth 2005),

*ρ* is the power parameter, restricted to the interval 1< *ρ* <2 (Dunn and Smyth 2005),

*λ*_*u*_ is a fixed effect for soak time categories,

δ_*n,ym*_ is a random effect to account for difference in landings resulting from spatiotemporal variation in fish density and/or catchability, according to NAFO subunit and year-month.

The offset accounts for the log of the number of gillnets used, relative to an assumed standard of 14 nets.

The estimated values *λ*_*u*_ were fitted to predicted values of landings from eq. 6, 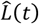 assuming log-normal error and the retention function was assumed to be 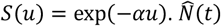 was calculated by solving the linear equation described earlier. All parameters for Model 1, including those of the GLMM, were estimated in a single negative log-likelihood function, thereby ensuring that uncertainty in estimated parameters and quantities best reflect uncertainties in the data.

#### 2.3.2 Model 2

The basic version of Model 2 was fitted to the data from field experiments exclusively. This version shared none of the data used in fitting Model 1, and therefore provides an estimate of catch losses that is completely independent. Two additional versions of Model 2 were fitted, incrementally adding data from additional sources.

Individual fish from the process study were classified as either dead-fresh or degraded, based on their condition upon retrieval, and grouped by individual net haul and whether they were attached to the net’s headline or leadline. The number of dead-fresh fish in haul *k* and on line *h*, given soak duration *u*_*k*_, *C*_*F,k,h*_(*u*_*k*_), was assumed to be a beta-binomial random variable,

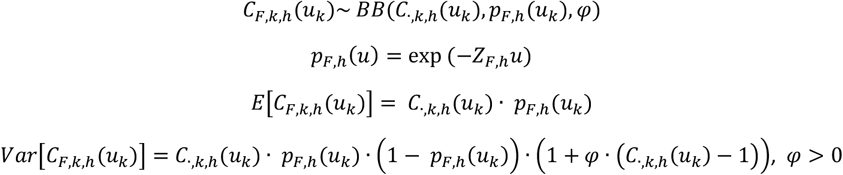

where,

*C*_·,*k,h*_(*u*_*k*_) is the total number of fish in process study haul *k* and line *h*,

*p*_*F,h*_(*u*) is the probability of being in the dead-fresh category for a fish attached to line *h* and after a soak duration *u*, and

*φ* is the beta-binomial dispersion parameter.

The beta-binomial assumption was used instead of the more common binomial assumption because of evidence of extra-binomial variation in the data. The above model estimates two degradation rate parameters, *Z*_*F,h*=*healine*_ and *Z*_*F,h*=*leadline*_ given that degradation was observed to occur more rapidly for fish tied to the leadline. In the absence of information on the degradation rate of the average Greenland halibut caught in a gillnet, we simply averaged these two values to estimate *Z*_*F*_.

The proportions of fish in each of the three observable catch condition categories (live, dead-fresh, degraded) were calculated for each experimental haul and were modelled using the multiplicative logistic normal distribution based on the continuation ratio logit (crl) transformation of the proportions. This model is particularly appropriate for ordered categories when a sequential process generates the response (Agresti 2002), and has been used to model age-compositions in stock assessment (Cadigan 2016). Indexing catch condition category as *c*, the crl proportions, *X*_*c,k*_, were computed as follows:

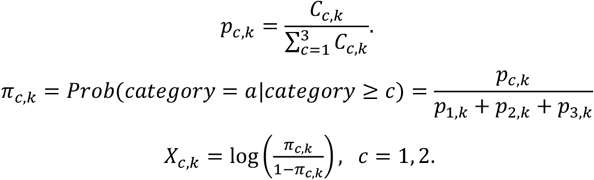

crl values *X*_*c,k*_ are obtained for both the observed and the model predicted catches. There are only two crl’s derived from the three catch proportions because those proportions sum to 1. The difference between observed and model predicted crl vectors was assumed to follow a normal distribution.

The landable catch from haul *k, C*_*live,k*_ + *C*_*dead*−*fresh,k*_ was fitted to the model predicted landings for a haul with the same soak duration, 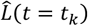 multiplied by an experiment-specific catchability parameter, *q*_*e*_, assuming log-normal error. The *q*_*e*_ parameters account for differences in available fish density during each experiment and are effectively nuisance parameters. The parameter value for the first experiment was set to 1 for model identifiability.

There were few observations for soak durations <18 hours in the field experiments, and all were exclusively from the fourth experiment. The basic fit for Model 2 therefore excluded data for soak duration <18 hours and as a consequence assumed that all the fish caught during the experiments were ‘regional’ fish, that is,

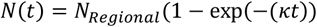

Subsequent variants excluded no data with respect to soak duration, and assumed the instantaneous net catch predicted by eq 7.

The second variant for Model 2 involved fitting additionally to the standardized catch rates as a function of soak duration from Chamberland and Benoît (2024). Attempts were made to fit to the original landings and effort data, as in the fitting of Model 1, but there were difficulties getting the model to converge. The availability of standardized catch information for soak duration <18 hours, in addition to observations available from the 4^th^ experiment, permitted the use of the complete instantaneous net catch function in eq 7.

The third variant for Model 3 involved fitting additionally to the catch condition data collected by at-sea observers.

## 3. Results

The estimated standardized catch rates for the fishery obtained by fitting Model 1 were very comparable to those obtained from the stand-alone estimation by Chamberland and Benoît (2024) using the GLMM, although the former were smoothed for longer soak durations (Figure 3a). This smoothing was expected given the smaller sample sizes and associated greater variability at longer soak duration, and the influence of the imposed model structure from eq. 6. Model 1, and the different variants of Model 2, fit the standardized catch rates well, with only small differences between models. One notable difference is that for Model 2 fit to all data including those from observers, which predicted an increase in catch in the first 3 hours of soak. Fits to the catches from the experiments were also good (Figure 3B). The fits to the catches in the 4^th^ experiment differed between the basic variant and the two others, because the former included only a subset of the data, resulting in a difference in the magnitude of the estimated catchability parameter.

Fish attached to the leadline in the process study degraded considerably more rapidly than those attached to the headline (Figure 4). There was also a notable difference in the degradation rate between experiments, with much slower rates in experiment 1, and rapid rates in experiment 4 leading to degradation of all individuals attached to the leadline within a couple hours.

**Figure 4.**
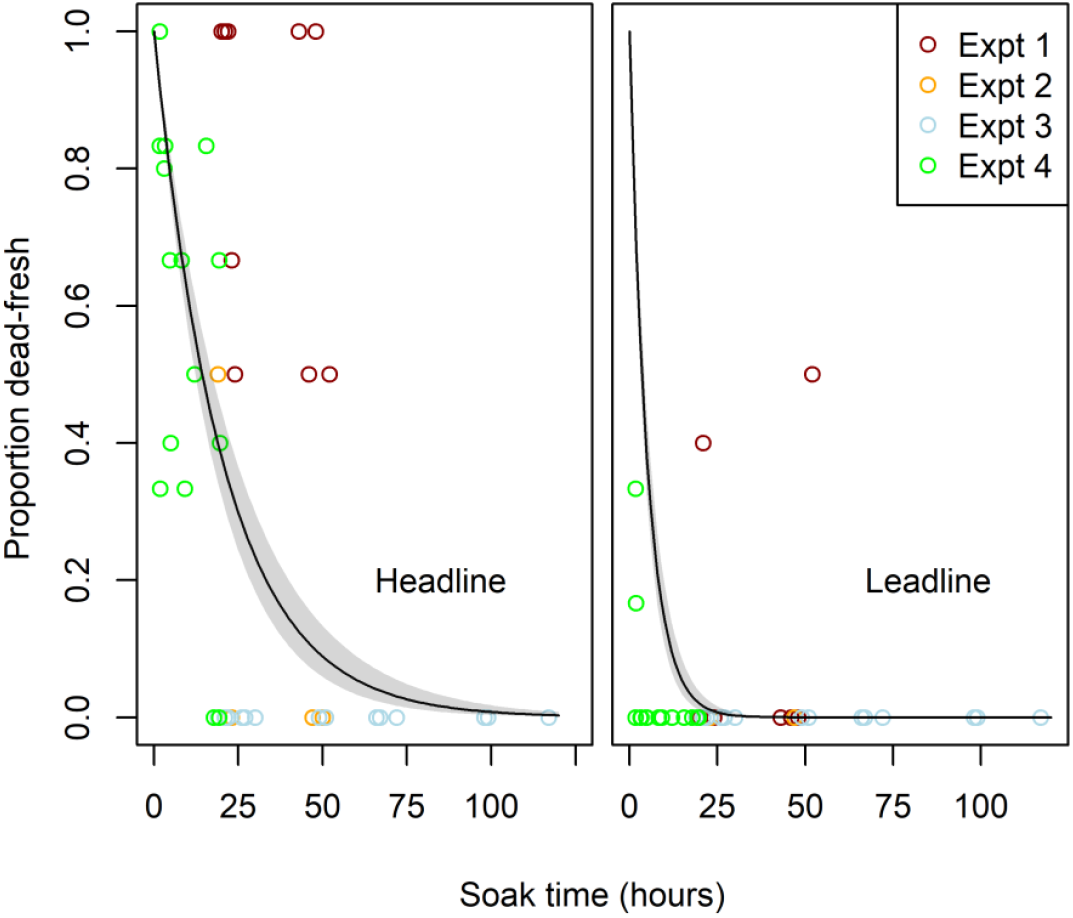
Results from the process study on the degradation rate of dead-fresh Greenland halibut attached to either the gillnet headline (left) or leadline (right). The proportion of fish in the dead-fresh category (as opposed to the degraded category) is plotted as a function of soak time duration (x axis) and experiment number (colours). The estimated degradation rate function and associated standard error are shown using the black line and shaded interval.

There was dispersion in the observed proportions of fish according to condition category, both within and between data source (Figure 5). Consistent with the results of the process study, for a given soak duration there was a higher proportion of live fish and lower proportion of degraded fish for experiment 1 compared to experiment 4. Results from at-sea observers, reflecting conditions in the broader fishery, were somewhat more aligned with those of experiment 1. Model selection using Akaike’s Information Criterion favored including the parameter for the fraction of long-term ‘hangers-on’, *p*_*d*_, for the basic fit of Model 2 (*p*_*d*_= 0.063 ± 0.027; ΔAIC = 4.85), but not the other two variants. This difference caused the basic fit for Model 2 to predict a decreasing proportion of live, and increasing proportion of degraded fish with soak duration, whereas the other variants predicted stable proportions for soak durations above about 30 hours, with values around 0.6 for live, 0.22 for dead-fresh and 0.18 for degraded fish (Figure 5). At shorter soak durations, the relevant Model 2 variants predict a rapid decrease in the proportion of live fish during the first 10 hours of soak, reaching a minimum around 18 hours, before increasing slightly to the asymptote values. Predictions for dead-fresh and degraded fish follow an opposite trend. Standardized crl residuals for the three variants of Model 2 do not display any notable trends over soak duration, but do reflect the experiment specific differences noted above (Figures S2-S4).

**Figure 5.**
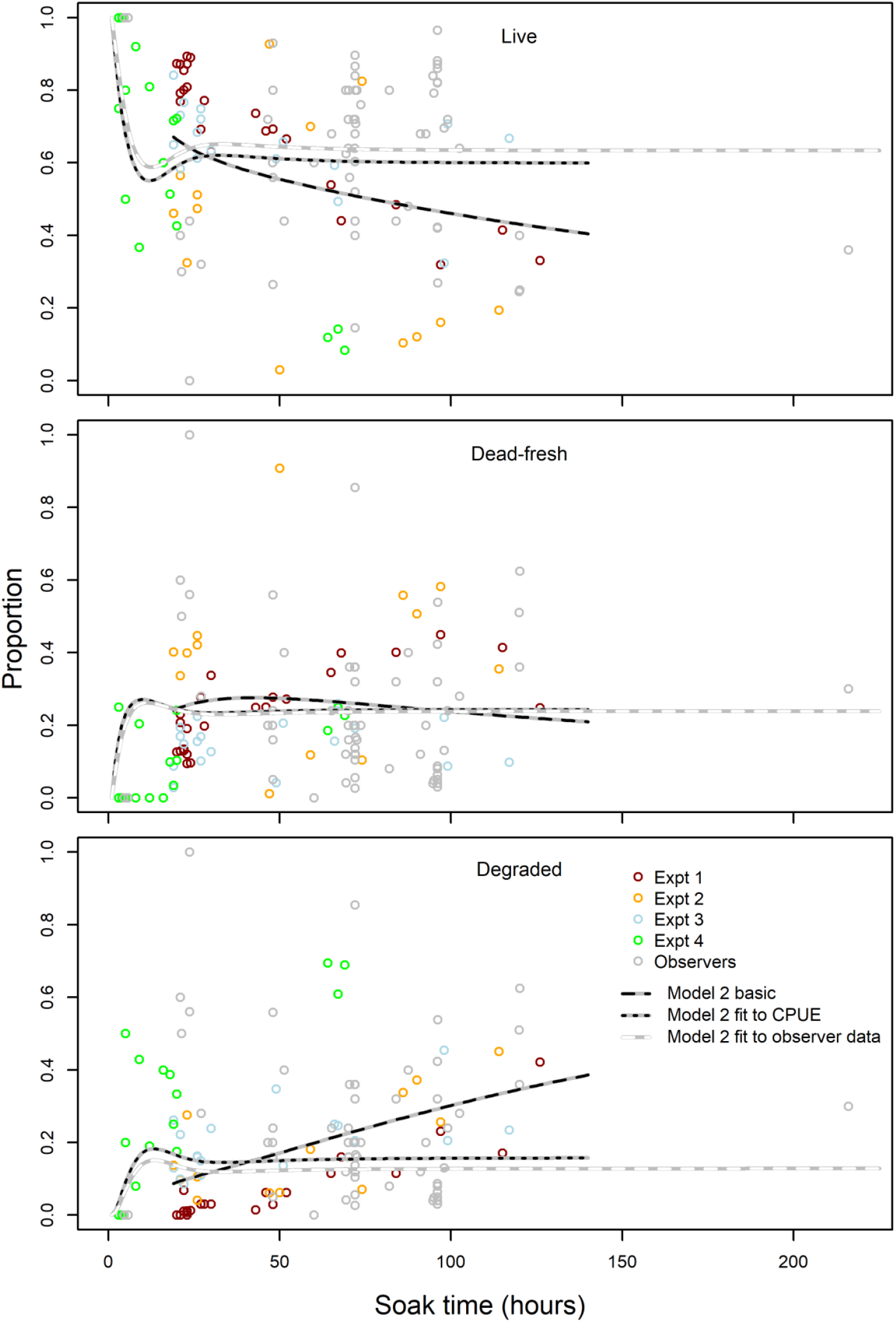
The proportion, by number, of Greenland halibut in each of the three condition categories (panels), as a function of soak duration (x axis) and by source, experiment number or at-sea observers (colours). The lines indicate the model estimated proportions for the three implementations of Model 2: fit to just the data from experiments (basic), fit to data from experiments and the estimated catch-per-unit-effort (CPUE), and fit to all data, including from at-sea observers.

The estimated retention function for Model 1 was more steeply declining that the functions estimated for Model 2 (Figure 6). The half-life of a fish in the net was about 6 hours for Model 1, and around 11.5 hours for Model 2. For both models, the proportion retained after 48 hours of soak was close to zero. The estimated dead-catch ratio functions for the models reflect the differences in predicted retention (Figure 7). The catch-ratio functions increase rapidly during the first 18 to 20 hours, reflecting the loss of the rapidly caught ‘local’ fish, and somewhat less rapidly thereafter. Overall, the models predict that the number of fish killed is twice the reported landings after ≤ 15 hours of soak for all models, about three times after 24 hours and close to 5 times after 72 hours, which is both the median and mode of soak durations in the fishery (Figure 1; Chamberland and Benoît 2024).

**Figure 6.**
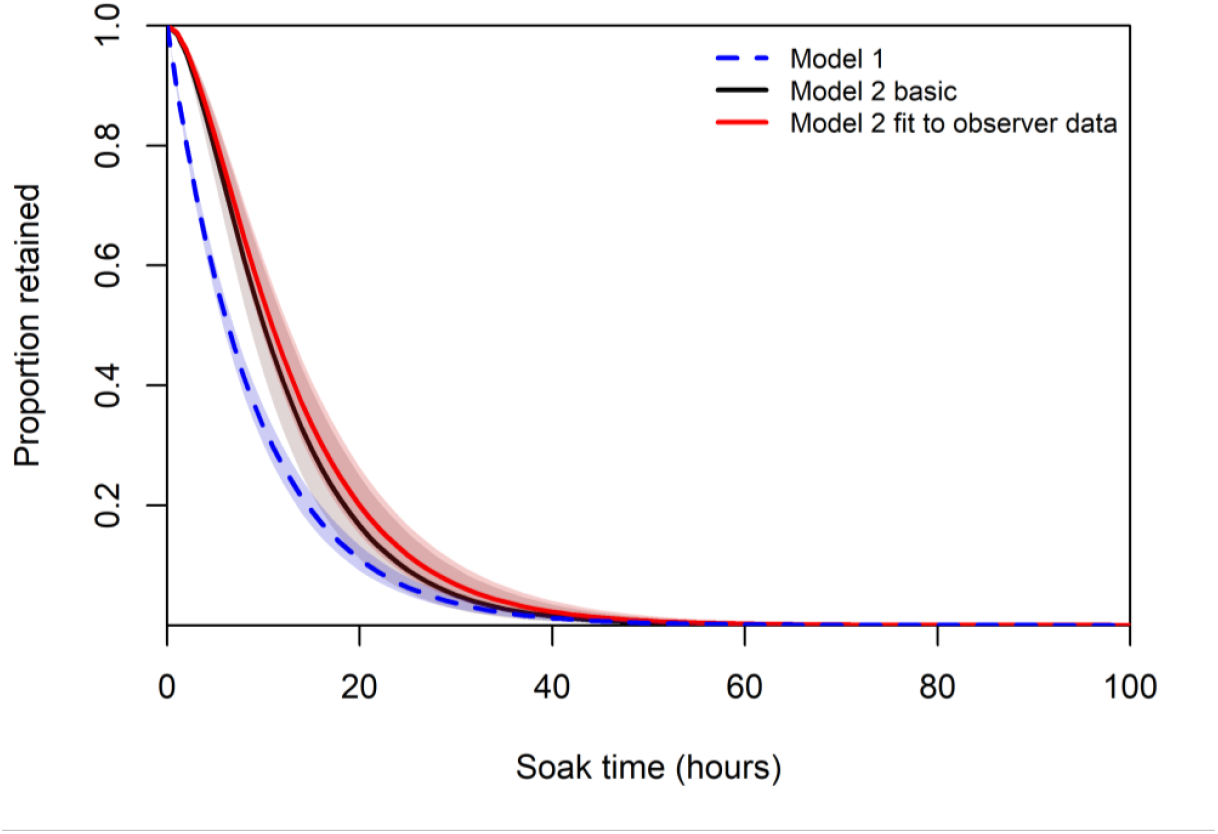
Estimated retention function with associated standard error (lines and shaded areas) for Model 1 and two implementations of Model 2, fit to just the data from experiments (basic) and fit to all data, including from at-sea observers. Given similarity in the results, only the predictions from the two most different versions of Model 2 are shown.

**Figure 7.**
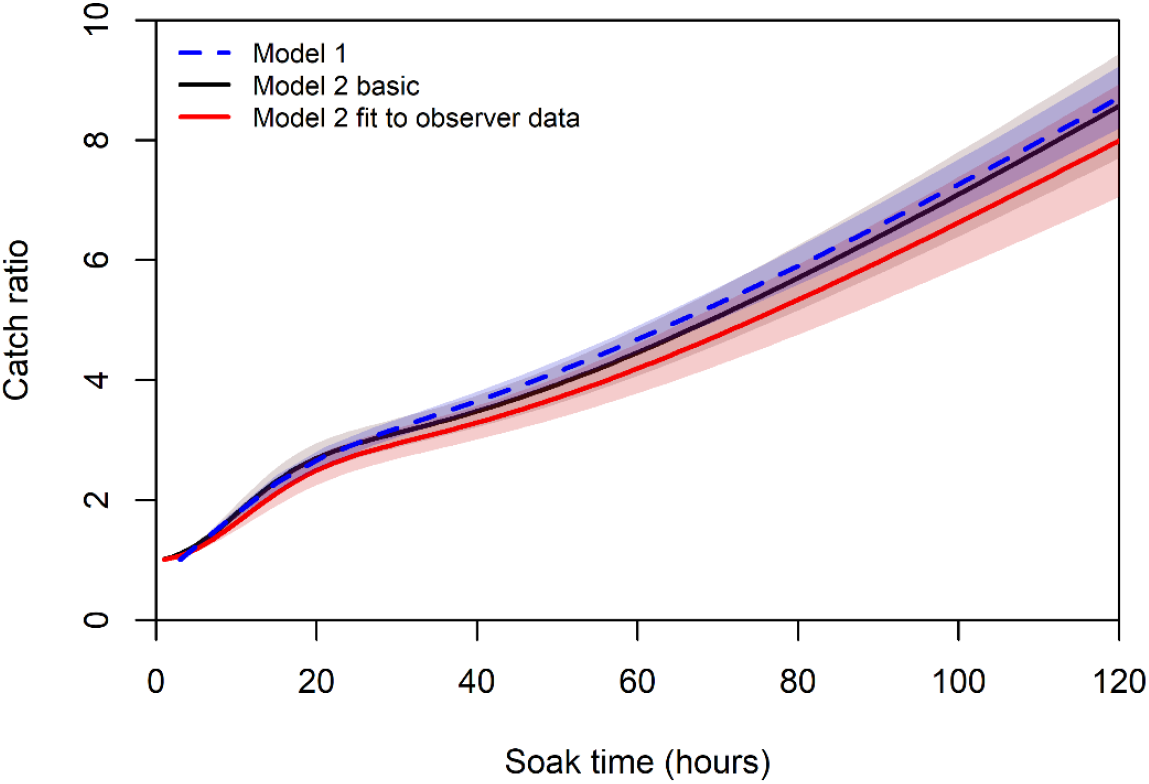
Estimated dead-catch ratio function *DC*(*t*_*i*_) with associated standard error (lines and shaded areas) for Model 1 and two implementations of Model 2, fit to just the data from experiments (basic) and fit to all data, including from at-sea observers. Given similarity in the results, only the predictions from the two most different versions of Model 2 are shown.

The annual mean fishery-scale dead-catch ratio differed by about 0.5 units between the higher predictions from Model 1 (mean 5.33 for 2000-2024) and the lower predictions from Model 2 fit to all data, including from observers (mean 4.78; Figure 8). The estimated annual mean ratios followed an oscillating trend, with peaks in 2005 and the late 2010s, and lower values around 2010 and more recently. Given that a single dead-catch ratio function was estimated for each model implementation, these trends reflect differences across years in soak durations employed in the fishery, and the retained catches that result from different soak times. The reasons for these interannual differences is presently not known. Nonetheless, the results show that even in years with a lower mean dead-catch ratio, a considerable amount of catch has been unaccounted-for.

**Figure 8.**
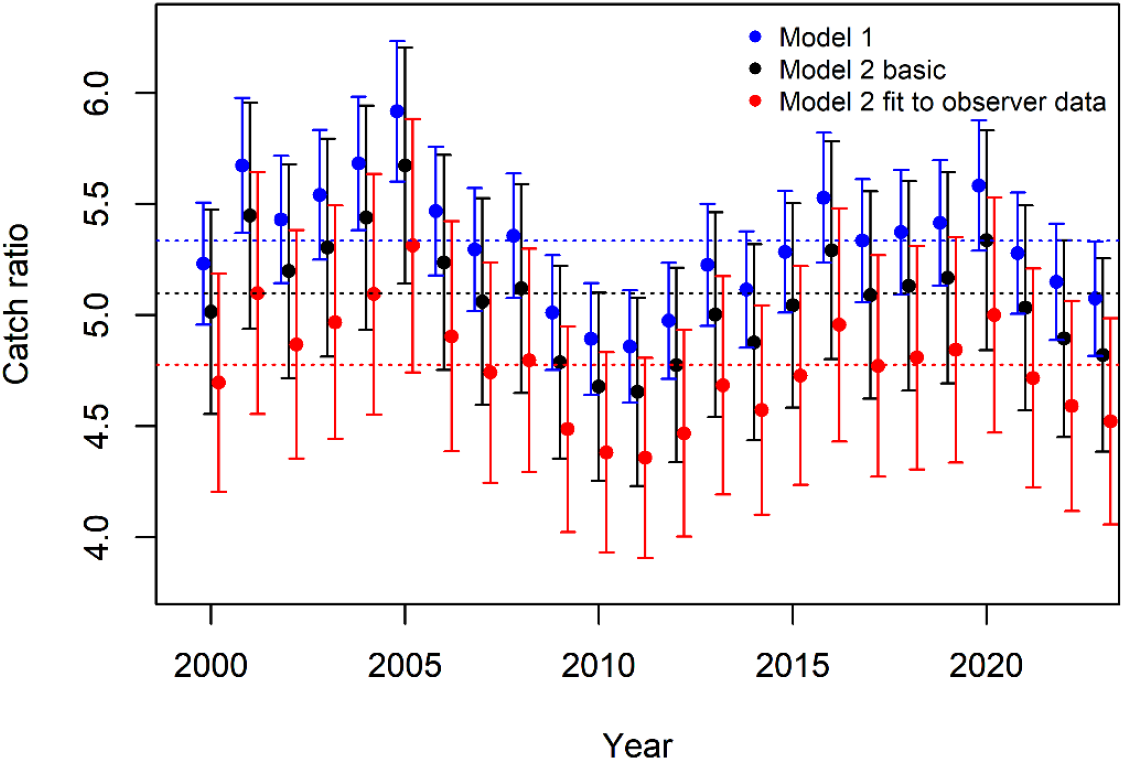
Estimated annual fishery-scale mean dead-catch ratio 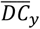 (with standard error) for Model 1 and two implementations of Model 2, fit to just the data from experiments (basic) and fit to all data, including from at-sea observers. Given similarity in the results, only the predictions from the two most different versions of Model 2 are shown.

## 4. Discussion

Globally, bottom-set gillnetting is the seventh most important fishing method by landed mass (derived from data provided in Pérez Roda et al. (2019)). This fishing method is also associated with the highest discarding rate (ratio of discarded and landed mass), after bottom-trawls and boat seines. While this likely reflects the somewhat unselective nature of gillnets which results in the catch of unwanted species (Kelleher 2005, Pérez Roda et al. 2019), the discarding of degraded or partially depredated catch is likely be an important contributor (Uhlmann and Broadhurst 2015, Tixier et al. 2021). For example, In a Portuguese hake (*Merluccius merluccius*) gillnet fishery, 65% of the catch was degraded and therefore discarded following 12 hours of soak time (Santos et al. 2002). Importantly, the results of the present study show that the rate at which target catch is discarded due to its degradation may belie considerable losses of dead fish. A median degraded catch discarding rate of about 5% after 72 hours of soaking (Figure S1) provided little a priori expectation of concurrent unaccounted dead catch losses estimated to be about four times the landings. Simply put, the state of catches when a gillnet is retrieved provides little indication of the turnover of fish prior to hauling.

The turnover of fish in a gillnet depends on the rate at which captured fish die, degrade and drop-out, and the capture of new catch, potentially attracted to the site. While mortality rates of gillnet caught fish are taxon, size and context specific (Uhlmann and Broadhurst 2015, Veldhuizen et al. 2018), they have been shown increase rapidly within minutes to a few hours following capture (e.g., Buchanan et al. 2002; Bell and Lyle 2016). For instance, in one experiment, 50% of coho salmon (*Oncorhynchus kisutch*) were dead within less than an hour following capture in a gillnet (Hargreaves and Tovey 2001). Similarly, the proportion of catch that is damaged or degraded can increase rapidly with soak time (e.g., Hickford and Schiel 1996; Savina et al. 2016). In the Portuguese hake fishery cited above, 23% of the catch was degraded after seven hours of soak time, almost tripling to 65% after 12 hours (Santos et al. 2002). Humborstad et al. (2003) found that individual fish could deteriorate completely within a 24-h period, due to the activity of fish lice in a deepwater Norwegian Greenland halibut fishery. This result is in line with the retention rate estimates of this study.

Our modelling approaches 1 and 2 (basic fit) estimate catch loss based on completely independent data sources. That these approaches estimate similar dead-catch ratios increases confidence in the estimates. The approaches share a common and fundamentally simple theoretical basis that seems conceptually reasonable to us and has the flexibility to estimate very small values of dead catch loss, if supported by the data. While the first approach infers turnover rates from strong signals in catch rates and may therefore be applicable only in a limited number of fisheries, the second approach estimated turnover rates directly from data that should be relative easily obtainable in most fishery contexts. Furthermore, our study has implications for studies of ghost fishing by abandoned, lost or discarded gillnets in which the condition of catches is examined over different gillnet soak durations (Humborstad et al. 2003). First and most obviously, given high similarity in experimental protocols, our second modelling approach should be easily transferable to these studies. Second, our result indicate clearly that fish losses resulting from decay can be substantially greater than might be implied by the composition of condition classes of the catch. The magnitude of losses from gillnet ghost fishing may have been substantially underestimated in previous studies (e.g., Kaiser et al. (1996);FAO (2016)) if turnover of fish in the net was not accurately quantified.

Results from the process study clearly indicate that degradation rates can be location and possibly time specific. This appears to be related to differences in the density of scavengers, evidenced by an abundance of amphipods (sand fleas) associated with catches during the fourth experience (J.-M. Chamberland, pers. obs.). This variability is further reflected in differences in the condition composition of catches with similar soak durations (Figure 5). Data were available for too few sites to properly characterize their contribution to variability in degradation rates, for example by estimating the degradation functions from a model including random effects for site and date. Such a model would likely better account for the variability observed in condition values between experiments and within the at-sea observer data. It would also likely improve the accuracy of the estimated degradation rates, and therefore the dead-catch ratios.

The magnitude of estimated annual mean dead-catch ratios for this fishery are considerable. Applied to annual recorded landings, the mean exploitation rate on the stock increases from about 6% to 30%, and the peak exploitation rate estimated for 1998 increases from about 16% to potentially 80% (Chamberland and Benoît 2024). The estimated dead-catch ratio series rests on the assumption that average catch retention rates are only a function of soak time. In addition to variability caused by local scavenger densities, it is likely that retention rates in the fishery have declined, and dead-catch ratios increased since 2010 as a result of an increase of about 1.5°C in the bottom waters occupied by GSL Greenland halibut (Galbraith et al. 2022, Chamberland and Benoît 2024). The rate of decay of dead fish by the action of microorganisms alone is a positive function of temperature (Groenewold and Fonds 2000). This is likely also the case for ectothermic scavengers such as hagfish and amphipods. While estimates from our first modelling approach are based on catch rates since 2000 and therefore likely represent average conditions over the period since then, estimates from our second approach are based on recent data and may therefore overestimate catch losses earlier in the period to an unknown degree.

### 4.1 Mitigating catch losses in the GSL fishery

Shortening gillnet soak times is the most effective means of reducing the unaccounted fishing mortality related to drop-outs. Result from the catch rate standardisation indicate that landable catches made with short soak duration (<6 hours) can be greater than those made with intermediate durations of 6-48 hours, and not much smaller than those employing longer soak durations that also effect considerably larger unaccounted catch amounts (Figure 3). Of course, the standard catch rates reflect average conditions for the 2000-2024 period, and not necessarily individual results. Some experimentation by harvesters employing different short soak durations is likely required to identify optimal durations. While these durations may yield lower catches that much longer ones, the ability to re-deploy the nets multiple times in a day, could result in overall higher retained catch and better quality of fish.

Estimated parameters for the catch rate model proposed in our the general framework, and results from the standardized catch rate analysis, suggest Greenland halibut local to the fishing site are rapidly captured and then subject to the degradation process. There is therefore some sensitivity as regards the best time to haul nets, which might not be fully characterized by the experimentation proposed above. A compromise for maintaining decent catches, while keeping soak times short, would be to fish formally baited nets as has been tried in the Greenland halibut fishery off Baffin Island, Canada (Bayse and Grant 2020). Baiting would increase the fishing power of the nets beginning as soon as they are deployed rather that having fishing power increase slowly as fish are first caught and then decay. However, the potential for increased catches of other predatory species, some of which may be of conservation concern, should be studied given the findings of Bayse and Grant (2020).

Apparent differences in degradation rates between experiments suggest that choosing sites with smaller densities of scavengers, should that be possible, could help mitigate drop-out losses. Fish harvesters may already posses this knowledge and there may be benefits to undertaking a questionnaire based survey of the industry to identify areas to avoid or favour. For some macrofaunal scavengers such as hagfish (*Myxine* sp.), data to identify fishing areas may already exist from trawl surveys (e.g., Morin et al. (2017)).

### 4.2 Conclusions

In this paper we present two modelling approaches for estimating the quantities of dead fish that drop out of gillnets, at least one of which should have broad applicability to other fisheries globally. The approaches produced very similar estimates of elevated catch losses that are consistent with the elevated mortality rates indicated by the stock assessment for GSL Greenland halibut. The results have broader implications for gillnet fisheries globally, including those employing relatively short soak durations. Notably, for relatively short soak times of ≤15 hours, the estimated magnitude of unobserved catch mortality was equivalent to that of the retained catch. Importantly, the physical condition of retained individuals may hide the true extent of fishing-induced mortality. Failure to incorporate such mortality into stock assessments may introduce substantial bias, potentially undermining the accuracy of population estimates and the robustness of sustainable harvest strategies. Additionally, these unaccounted losses represent a significant source of fishing-related waste.

## 5. Acknowledgements

Jun Allard provided advice on the mathematical approach used in the general modelling framework. We thank the captains, D. Cassivi, P.-A. Huard and P. N. Tanguay-Lévesque, and their crew, along with M.-C. Marquis, M.-M. Rondeau and G. Cortial for making the field experiments possible. We also thank the coordinator and at-sea observers at Biorex for accommodating our request to collect additional data on the condition of Greenland halibut during the commercial fishery. This study was funded under Fisheries and Oceans Canada’s Fisheries Science Collaborative Program.

## Appendix I.

### Supplementary Figures

**Figure S1.**
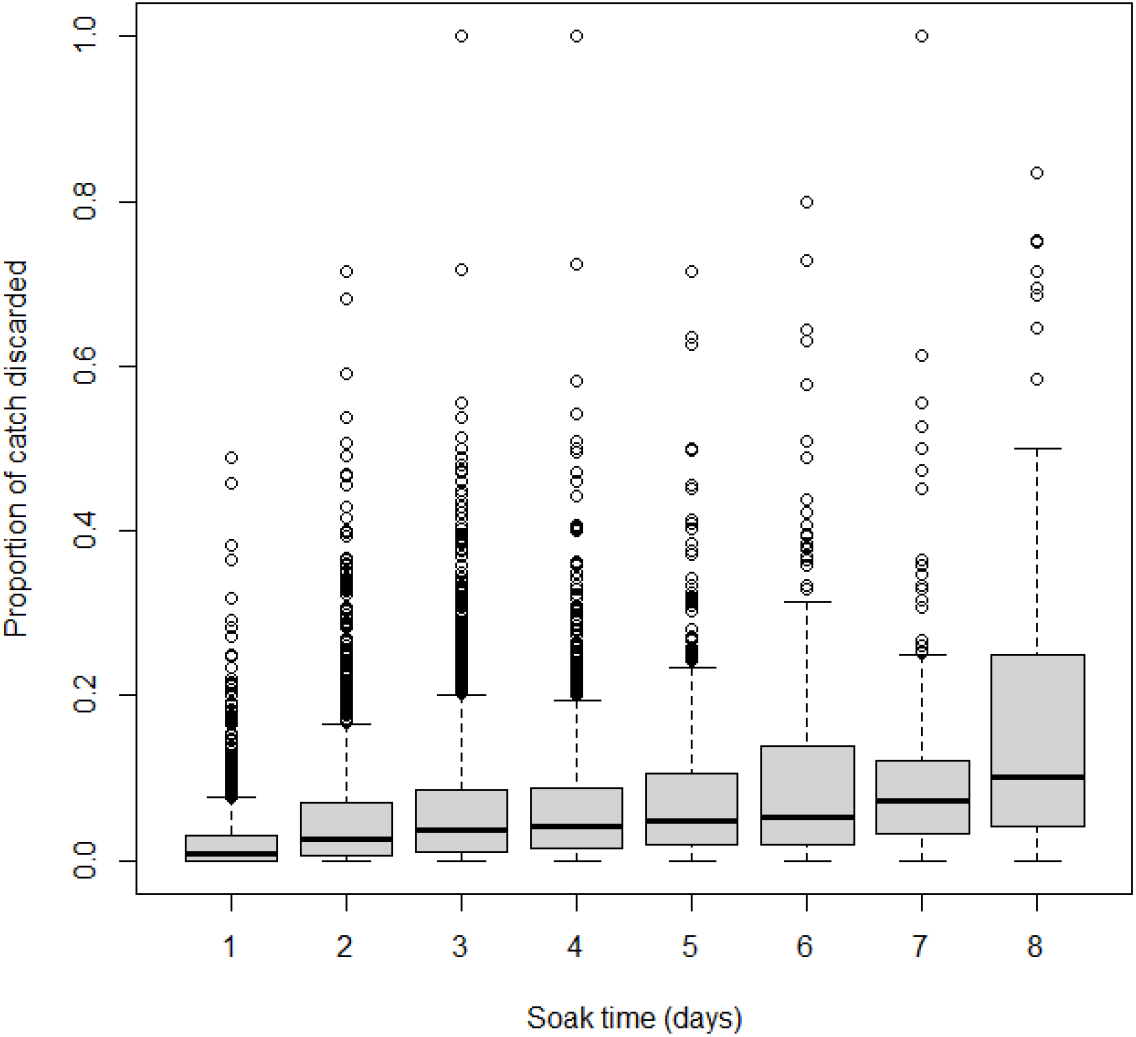
Boxplots of the proportion, by weight, of Greenland halibut catch discarded in the directed fishery as a function of soak duration (times rounded to integer days) from the at-sea observer data, 2000-2022. Note that category 1 is for ≤1 day and 8 is for *≥*8 days.

**Figure S2.**
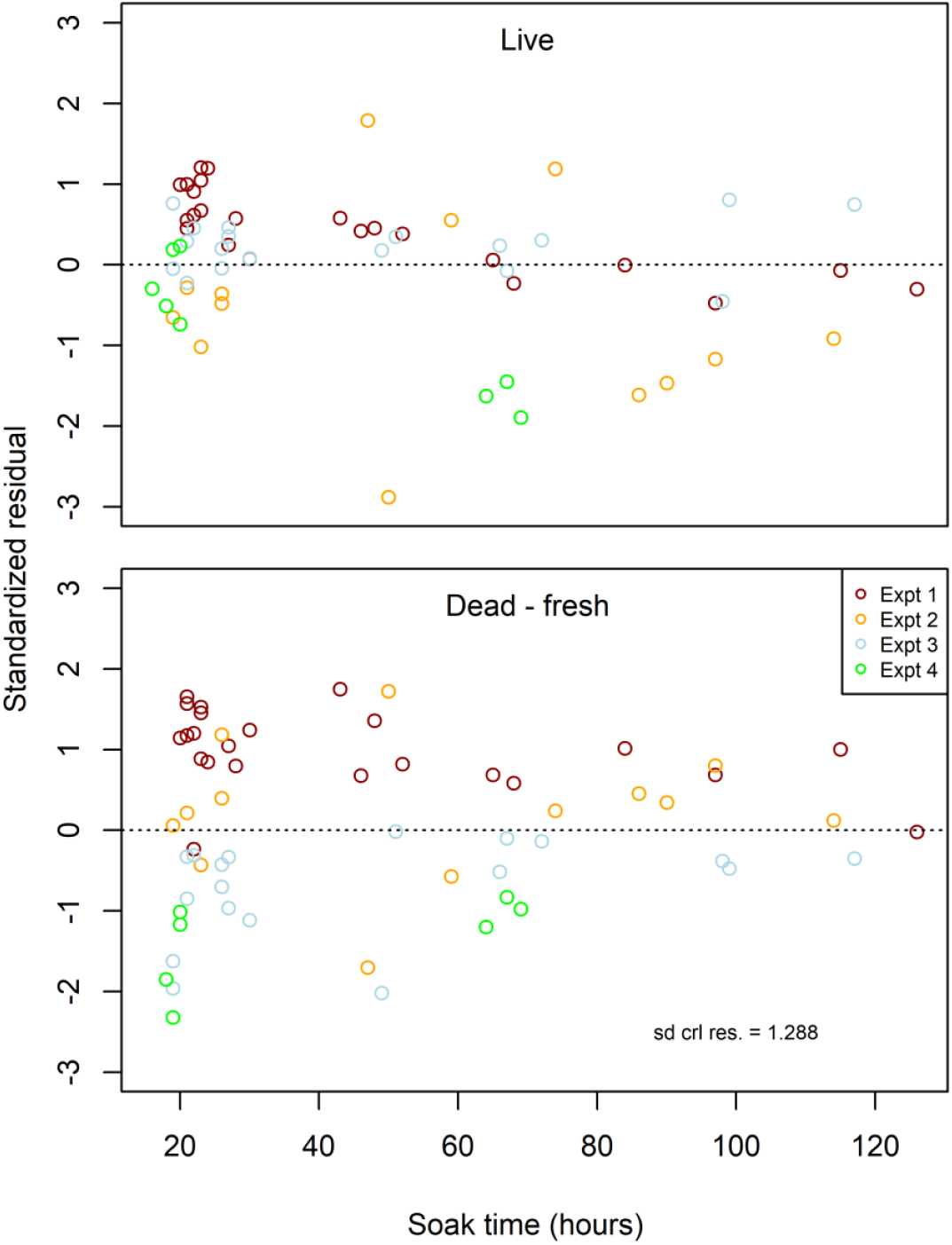
Standardized continuation ratio logit residuals for Model 2, fit to just the data from experiments (basic).

**Figure S3.**
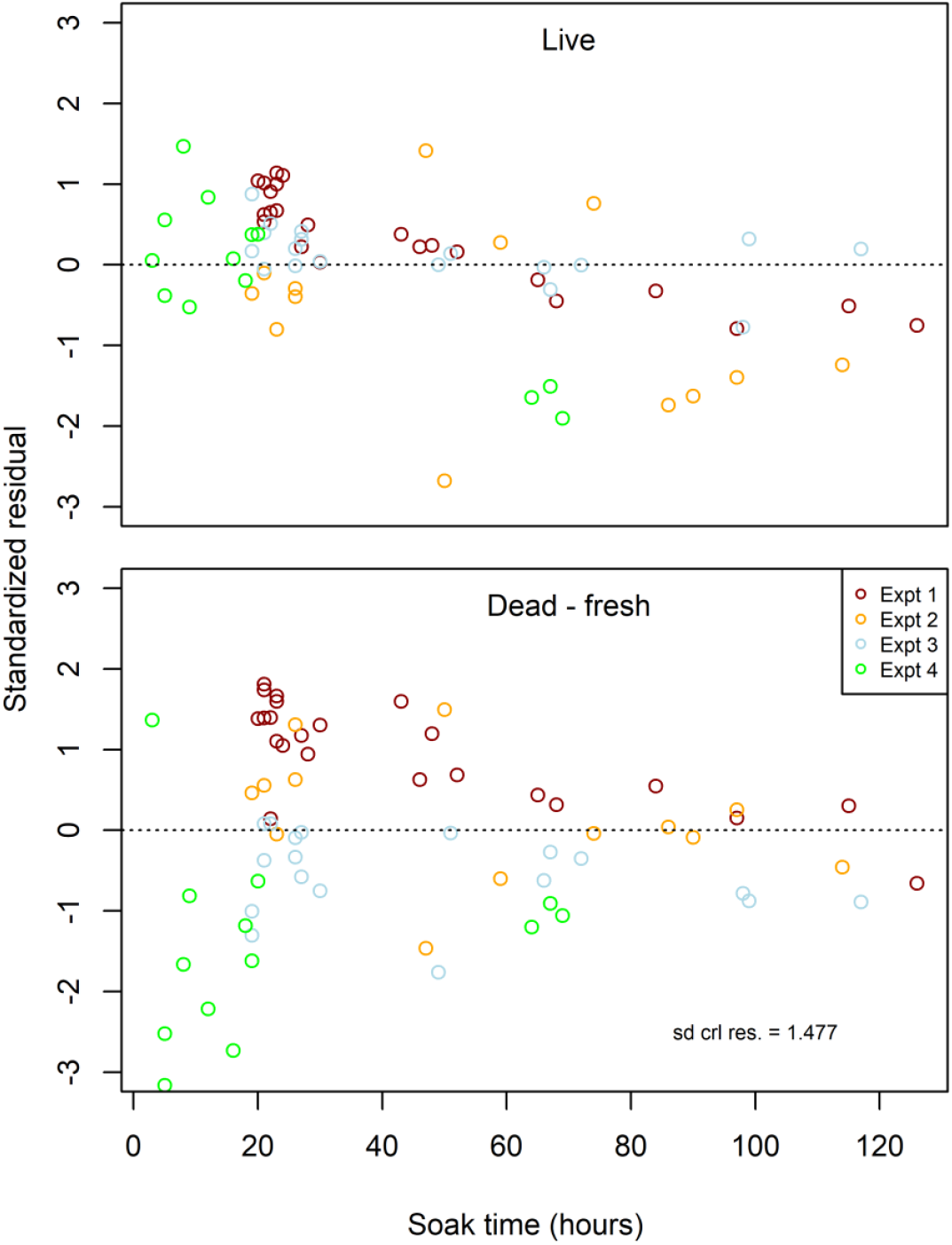
Standardized continuation ratio logit residuals for Model 2, fit to the data from experiments and the catch-per-unit-effort from the fishery.

**Figure S4.**
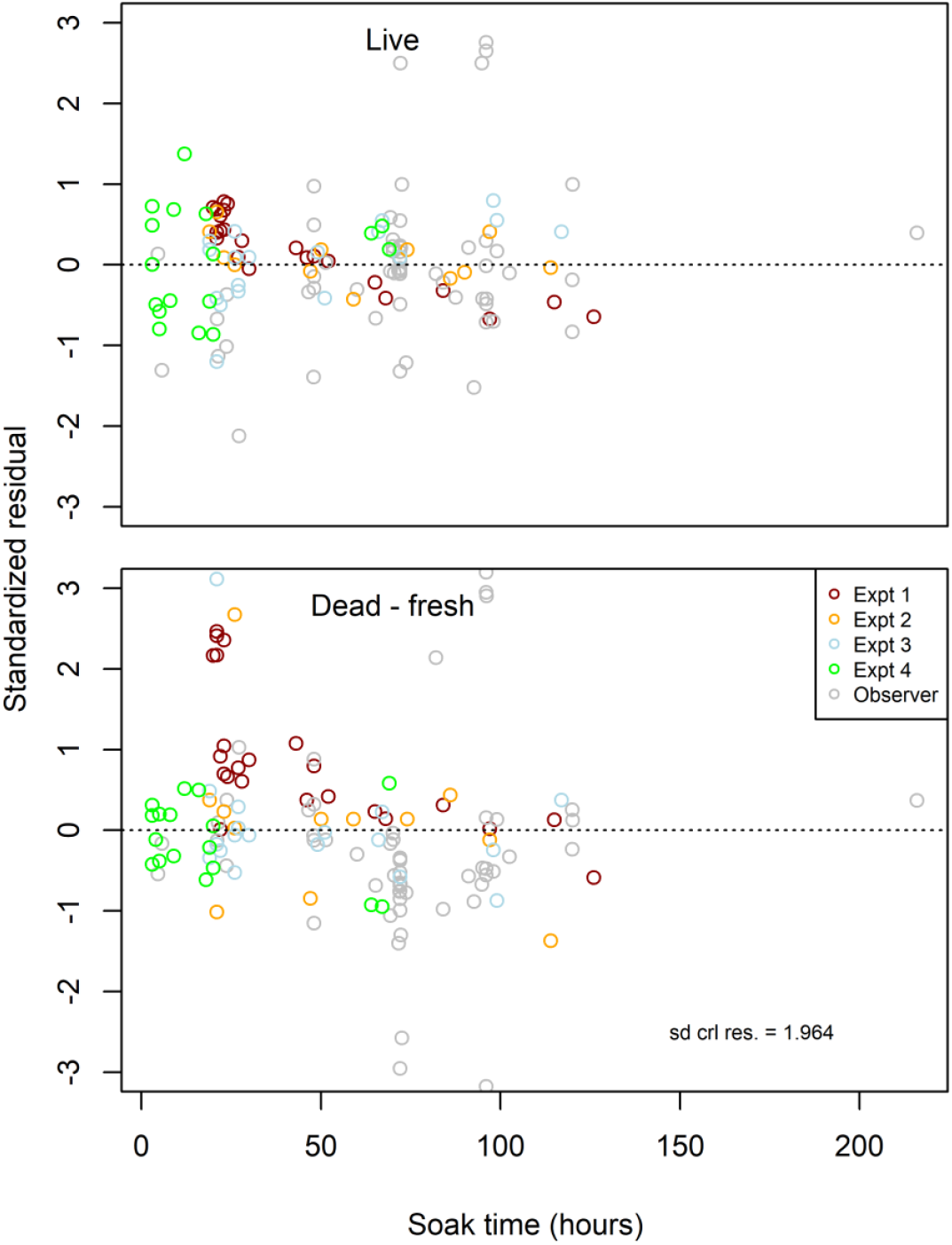
Standardized continuation ratio logit residuals for Model 2, fit to all the data, including from at-sea observers.

## Literature cited

Agresti, A. 2002. Categorical Data Analysis, 2nd ed. John Wiley and Sons Inc., New Jersey.

Bayse, S. M., and S. M. Grant. 2020. Effect of baiting gillnets in the Canadian Greenland halibut fishery. Fisheries Management and Ecology 27:523–530.

Bousquet, N., N. Cadigan, T. Duchesne, and L.-P. Rivest. 2010. Detecting and correcting underreported catches in fish stock assessment: trial of a new method. Canadian Journal of Fisheries and Aquatic Sciences 67:1247–1261.

Brogan, J. D., C. R. Kastelle, T. E. Helser, and D. M. Anderl. 2021. Bomb-produced radiocarbon age validation of Greenland halibut (Reinhardtius hippoglossoides) suggests a new maximum longevity. Fisheries Research 241:106000.

Cadigan, N. G. 2016. A state-space stock assessment model for northern cod, including under-reported catches and variable natural mortality rates. Can. J. Fish. Aquat. Sci. 73:296–308.

Chamberland, J.-M., and H. Benoît. 2024. Gulf of St. Lawrence (4RST) Greenland Halibut stock status in 2022. DFO Can. Sci. Advis. Sec. Res. Doc. 2024/001. v + 144 p.

Cook, R. M. 2019. Inclusion of discards in stock assessment models. Fish and Fisheries 20:1232–1245.

Dickey-Collas, M., M. A. Pastoors, and O. A. van Keeken. 2007. Precisely wrong or vaguely right: simulations of noisy discard data and trends in fishing effort being included in the stock assessment of North Sea plaice. ICES Journal of Marine Science 64:1641–1649.

Dunn, P. K., and G. K. Smyth. 2005. Series evaluation of Tweedie exponential dispersion model densities.. Stat. Comput. 15:267–280.

Engås, A., T. Jørgensen, and K. K. Angelsen. 2000. Effects on catch rates of baiting gillnets. Fisheries Research 45:265–270.

FAO. 2016. Abandoned, lost and discarded gillnets and trammel nets: methods to estimate ghost fishing mortality, and the status of regional monitoring and management, by Eric Gilman, Francis Chopin, Petri Suuronen and Blaise Kuemlangan.

Galbraith, P. S., J. Chassé, J. Dumas, J.-L. Shaw, C. Caverhill, D. Lefaivre, and C. Lafleur. 2022. Physical Oceanographic Conditions in the Gulf of St. Lawrence during 2021. DFO Can. Sci. Advis. Sec. Res. Doc. 2022/034. iv + 83 p.

Glemarec, G., A.-M. Kroner, and L. Kindt-Larsen. 2024. Disappearing fish: Grey seal depredation in a Baltic net fishery. Fisheries Research 277:107070.

Groenewold, S., and M. Fonds. 2000. Effects on benthic scavengers of discards and damaged benthos produced by the beam-trawl fishery in the southern North Sea. ICES Journal of Marine Science 57:1395–1406.

Hargreaves, N. B., and C. Tovey. 2001. Mortality rates of coho salmon caught by commercial salmon gillnets and the effectiveness of revival tanks and reduced soak time for decreasing coho mortality rates. DFO Can. Sci. Advis. Sec. Res. Doc. 2001/154:56 p.

Humborstad, O.-B., S. Løkkeborg, N.-R. Hareide, and D. M. Furevik. 2003. Catches of Greenland halibut (Reinhardtius hippoglossoides) in deepwater ghost-fishing gillnets on the Norwegian continental slope. Fisheries Research 64:163–170.

ICES. 2005. Joint report of the study group on unaccounted fishing mortality (SGUFM) and the workshop on unaccounted fishing mortality (WKUFM).. ICES CM 2005/B:08, 68 pp.

Kaiser, M., B. Bullimore, P. Newman, K. Lock, and S. Gilbert. 1996. Catches in ‘ghost fishing’ set nets. Mar. Ecol. Prog. Ser. 145:11–16.

Kelleher, K. 2005. Discards in the worlds fisheries: an update.. Food and Agriculture Organisation Fisheries Technical Paper, 470, Food and Agriculture Organisation Fisheries department, Rome, Italy.

Kristensen, K. 2024. RTMB: ‘R’ Bindings for ‘TMB’. R package version 1.6. https://github.com/kaskr/rtmb.

Maunder, M. N., and A. E. Punt. 2004. Standardizing catch and effort data: a review of recent approaches. Fisheries Research 70:141–159.

Morin, R., D. Ricard, H. Benoît, and T. Surette. 2017. A review of the biology of Atlantic hagfish (Myxine glutinosa), its ecology, and its exploratory fishery in the southern Gulf of St. Lawrence (NAFO Div. 4T). DFO Can. Sci. Advis. Sec. Res. Doc. 2017/017. v + 39 p.

Patterson, D. A., K. A. Robinson, R. J. Lennox, T. L. Nettles, L. A. Donaldson, E. J. Eliason, G. D. Raby, J. M. Chapman, K. V. Cook, M. R. Donaldson, A. L. Bass, S. M. Drenner, A. J. Reid, S. J. Cooke, and S. G. Hinch. 2017. Review and evaluation of fishing-related incidental mortality for Pacific salmon. DFO Can. Sci. Advis. Sec. Res. Doc. 2017/010. ix +155 p.

Pérez Roda, M., E. Gilman, T. Huntington, S. Kennelly, P. Suuronen, M. Chaloupka, and P. Medley. 2019. A third assessment of global marine fisheries discards.

Perretti, C. T., J. J. Deroba, and C. M. Legault. 2020. Simulation testing methods for estimating misreported catch in a state-space stock assessment model. ICES Journal of Marine Science 77:911–920.

Punt, A. E., D. C. Smith, G. N. Tuck, and R. D. Methot. 2006. Including discard data in fisheries stock assessments: Two case studies from south-eastern Australia. Fisheries Research 79:239–250.

Santos, M., M. Gaspar, C. C. Monteiro, and P. Vasconcelos. 2002. Gill net and long-line catch comparisons in a hake fishery: The case of southern Portugal. Scientia Marina 66:433–441.

Tixier, P., M.-A. Lea, M. A. Hindell, D. Welsford, C. Mazé, S. Gourguet, and J. P. Y. Arnould. 2021. When large marine predators feed on fisheries catches: Global patterns of the depredation conflict and directions for coexistence. Fish and Fisheries 22:31–53.

Uhlmann, S. S., and M. K. Broadhurst. 2015. Mitigating unaccounted fishing mortality from gillnets and traps. Fish and Fisheries 16:183–229.

Veldhuizen, L. J. L., P. B. M. Berentsen, I. J. M. de Boer, J. W. van de Vis, and E. A. M. Bokkers. 2018. Fish welfare in capture fisheries: A review of injuries and mortality. Fisheries Research 204:41–48.

Ward, P., R. A. Myers, and W. Blanchard. 2004. Fish lost at sea: the effect of soak time on pelagic longline catches. Fishery Bulletin 102:179–195.

